# Spatial patterns of mosquito communities and monkey malaria vectors in a tropical riverine forest in Sabah, Malaysian Borneo

**DOI:** 10.64898/2026.04.27.720977

**Authors:** Ikki Matsuda, Benny Obrain Manin, Takaaki Yahiro, Primus Lambut, Joseph Tangah, Michael A. Huffman, Henry Bernard, Vijay Subbiah Kumar, Tock Hing Chua

**Author notes:** Correspondence: Tock Hing Chua & Ikki Matsuda.

## Abstract

**Background:** Understanding how vector ecology intersects with host behaviour is essential for predicting zoonotic malaria risk in tropical ecosystems. In Malaysian Borneo, zoonotic transmission of simian malaria parasites such as *Plasmodium knowlesi* is shaped by interactions among mosquito vectors, non-human primate reservoir hosts, and human activities across heterogeneous landscapes. However, fine-scale spatial structuring of mosquito communities within tropical riverine forests remains poorly characterized.

**Methodology/Principal Findings:** We conducted a two-year mosquito survey (November 2016–October 2018) in a lowland riverine forest along the Menanggul River, Sabah, Malaysian Borneo. Mosquitoes were sampled on 44 nights using CO₂-baited light traps repeatedly deployed across horizontal (0 m, 250 m, and 500 m from the river) and vertical (ground and canopy) gradients, yielding 244 trap collections. A total of 9,928 mosquitoes were collected, dominated by *Culex* spp. (91.4%), whereas *Anopheles* spp. were rare (153 individuals; 1.5%). Female *Anopheles* mosquitoes (n = 57) were more frequently detected near the river and tended to occur more often in ground traps. Zero-inflated negative binomial generalized linear mixed models based on the female mosquito dataset revealed significant effects of river distance and trap height on mosquito abundance, while short-term microclimatic variation showed little explanatory power. The *Anopheles* assemblage was dominated by *Anopheles balabacensis*, a recognized vector of simian malaria parasites in Sabah. Molecular screening detected simian *Plasmodium* DNA in two *An. balabacensis* individuals, including *P. knowlesi*, together with other simian *Plasmodium* species.

**Conclusions/Significance:** Our findings suggest that river-edge habitats may function as localized contact hotspots where mosquito vectors, primate reservoir hosts, and human activities repeatedly overlap in space and time, potentially facilitating zoonotic malaria transmission even when vector densities are low. These results support the growing view that zoonotic malaria risk in tropical landscapes is shaped less by overall vector abundance than by fine-scale ecological overlap among vectors, wildlife, and humans.

**Author Summary:** Zoonotic malaria caused by monkey malaria parasites such as *Plasmodium knowlesi* has become an important public health concern in Malaysian Borneo. Transmission occurs when mosquito vectors feed on both infected non-human primates and humans, but the ecological conditions that increase opportunities for such contact remain poorly understood. In this study, we investigated mosquito communities in a tropical riverine forest along the Menanggul River in Sabah, Malaysia, an area where wildlife, human activities, and ecotourism frequently overlap. Over a two-year period, we repeatedly sampled mosquitoes across different distances from the river and at both ground and canopy levels. Although mosquitoes capable of transmitting simian malaria were relatively rare, they were detected more frequently near river edges and in ground-level traps. We also detected simian *Plasmodium* DNA in two *Anopheles balabacensis* mosquitoes, a major vector of zoonotic malaria in Sabah. Our findings suggest that river-edge habitats may act as localized “contact hotspots” where vectors, primate hosts, and humans repeatedly overlap in space and time, potentially increasing opportunities for zoonotic malaria transmission even when overall vector abundance is low. These results highlight the importance of considering wildlife ecology, human behaviour, and landscape structure together when assessing and managing zoonotic disease risk in tropical forest systems.

## Introduction

Mosquitoes play a central role in the transmission of vector-borne diseases, and understanding the ecological factors that shape mosquito abundance and distribution is critical for predicting disease risk in tropical ecosystems (1, 2). In particular, malaria transmission is determined not only by the presence of competent vectors, but also by when and where vectors and hosts co-occur, together with environmental conditions that shape vector and parasite traits relevant to transmission (3–5). While climatic variables such as temperature and humidity can influence mosquito development and survivorship (6, 7), the distribution of adult mosquitoes is often strongly structured by habitat features, especially the availability and spatial configuration of aquatic breeding sites (2, 8). In forested environments, mosquito communities may also exhibit vertical stratification, reflecting variation in host availability and microhabitat conditions across canopy and ground strata (9).

In Southeast Asia, these ecological processes, especially those modified by anthropogenic land-use changes, are closely linked to the emergence of zoonotic malaria. In particular, infections caused by *Plasmodium knowlesi*, a simian malaria parasite, have become a major public health concern in Malaysian Borneo (10–12). Transmission of simian malaria involves complex interactions among mosquito vectors, non-human primate reservoir hosts, and human populations. In Malaysian Borneo, several *Anopheles* species in the Leucosphyrus Group, especially *Anopheles balabacensis*, are recognized as primary vectors of simian malaria parasites (13–15). Human-landing studies have shown that *A. balabacensis* preferentially bites at ground level while still utilizing canopy strata, supporting its role as a potential bridge vector between arboreal primates and ground-dwelling humans in disturbed forest systems (9). This vertical flexibility suggests that relatively small shifts in host use of canopy versus ground habitats could influence opportunities for zoonotic transmission in forest-edge environments. In addition, increasing evidence suggests that vector abundance alone does not adequately predict transmission risk. Instead, risk is shaped by the spatial overlap among vectors, reservoir hosts, and human activities across heterogeneous landscapes, including forest, agricultural, and village environments (16–18).

A growing number of studies in Sabah, Bornean Malaysia, have examined how land use, environmental conditions, and human behaviour influence exposure to simian malaria vectors. For example, vector abundance and species composition vary across gradients of forest disturbance and land use (19, 20), and human exposure to zoonotic-malaria vectors increases when people spend time in forest and forest-edge settings (e.g. farming, forest product collection, or other outdoor activities), where contact with host-seeking mosquitoes is more likely (16, 17, 21). Complementary landscape-scale analyses further show that forest fragmentation, proximity to plantation agriculture, and clearing around households are strongly associated with the distribution of *P. knowlesi* vector habitats and human exposure (20, 22). These findings indicate that anthropogenic land-use change reshapes vector ecology and creates new transmission interfaces. Within this context, riverine forests may function as ecological linkages connecting disturbed forests, plantation landscapes, and human settlements, yet mosquito community structure within these systems remains poorly characterized.

In addition, studies focusing on primate hosts have shown that macaques serve as important reservoir hosts for simian malaria parasites (23, 24), and that vector-host interactions can vary across habitat types and vertical strata (9, 24). Recent work in Sabah further indicates that *A. balabacensis* can sustain *P. knowlesi* transmission even at relatively low infection rates and vector densities, as biting activity is highly concentrated in specific habitats and times of day (14, 20, 25, 26). This suggests that fine-scale spatial and temporal overlap among vectors, primate hosts, and humans, rather than overall vector abundance, may be a key determinant of zoonotic malaria risk in fragmented Bornean landscapes. Despite these advances, fine-scale spatial patterns of mosquito communities within riverine forest systems remain poorly understood, particularly in relation to vertical stratification and proximity to aquatic habitats.

Riverine forests form key ecological interfaces in tropical landscapes, where aquatic habitats, wildlife activity, and human use often converge. In the Lower Kinabatangan region of Sabah, river-edge habitats support diverse primate assemblages and are also focal areas for human activities such as ecotourism (27, 28). Many primate species, including macaques and proboscis monkeys, utilize river margins particularly during dawn and dusk, when they are most readily observed from boats (29–31). These predictable patterns of wildlife and human activity make river margins a logical setting in which to examine how mosquito communities, including potential vectors, are distributed across space.

Despite the potential importance of riverine habitats in shaping vector-host interactions, few studies have simultaneously examined mosquito community structure, vertical stratification, and distance from aquatic habitats in a single framework (14–16, 22, 26). In particular, the extent to which potential simian malaria vectors exhibit spatial structuring across riverine gradients and forest strata remains unclear. Moreover, it is not well understood whether short-term microclimatic variation provides additional explanatory power beyond spatial factors in shaping mosquito abundance in these systems. In this study, therefore, we conducted systematic mosquito sampling in a tropical riverine forest along the Menanggul River, a tributary of the Kinabatangan River in Sabah, Malaysian Borneo. Using standardized light- and CO₂-baited traps deployed across horizontal (distance from river) and vertical (ground vs canopy) gradients, we aimed to (1) characterize mosquito community composition, (2) quantify spatial variation in mosquito abundance, (3) examine the distribution of potential simian malaria vectors (*Anopheles* spp.) and (4) assess the relative importance of spatial versus microclimatic factors in shaping mosquito abundance. In addition, we screened *Anopheles* mosquitoes for *Plasmodium* infections to evaluate the presence of simian malaria parasites within the study area. Finally, by integrating community-level patterns with spatial and epidemiological perspectives, we attempted to provide insights into how fine-scale ecological structure shapes vector distribution and potential zoonotic risk in tropical forest ecosystems.

## Methods

### Study site

This study was conducted in lowland riverine forests along the Menanggul River, a tributary of the Kinabatangan River, Sabah, Malaysia (118° 30ʹ E, 5° 8ʹ N). The southern area of the Menanggul River is extensively covered by secondary forest, whereas the northern area has been deforested for oil palm plantations, except for a protected zone along the river (Supplementary Figure S1). Regionally, the mean minimum and maximum daily temperatures were approximately 24 °C and 30°C, and the mean annual precipitation at the site was 2,474 mm (32). The study site supports several sympatric primate species, including proboscis monkeys (*Nasalis larvatus*), long-tailed macaques (*Macaca fascicularis*), pig-tailed macaques (*M. nemestrina*), silvered langurs (*Trachypithecus cristatus*), maroon langurs (*Presbytis rubicunda*), Hose’s langurs (*P. hosei*), Bornean gibbons (*Hylobates muelleri*) and orangutans (*Pongo pygmaeus*) (29). Long-tailed macaques and pig-tailed macaques are the recognized major natural hosts of *P. knowlesi* (33). The two *Presbytis* spp. are potential candidates at the study site, given that infection has been reported in a closely related species, *P. melalophos*, in Peninsular Malaysia and Indonesia (34).

### Mosquito collection with environmental measurements

Mosquito collections were carried out from November 2016 to October 2018 at a frequency ranging from one to three times per month. On each trapping date, we deployed up to six traps simultaneously. Mosquito trapping was conducted on 44 dates. Of the planned 264 trap-nights (44 dates × 6 trap positions), 244 trap collections (92.4%) were successfully completed. The remaining 20 trap-nights were lost because seasonal flooding prevented access to the inland trapping locations (250 m and 500 m), whereas both river-edge (0 m) traps were successfully sampled on all trapping dates. Mosquitoes were captured using a fan-based, battery-operated vacuum trap equipped with a miniature light bulb and a carbon dioxide source generated by mixing commercial yeast and sugar water in a 50 ml tube (Figure 1). Light- and CO₂-baited fan traps are widely used to index host-seeking mosquito activity; however, their efficiency varies among taxa and may not directly approximate human landing rates, particularly for *Anopheles* vectors. Studies in Sabah have shown that human landing catches can yield different estimates of *Anopheles* abundance compared to trap-based collections, highlighting that trap-based estimates may underestimate realized human biting exposure (16). To obtain environmental data, temperature and humidity loggers were attached to each trap and recorded at hourly intervals. Six trap locations were defined along a 500-m transect extending perpendicularly inland from the right bank of the Menanggul River, approximately 3500 m upstream from the river mouth. Traps were installed at two vertical strata: 1.5 m above ground level (“ground traps”) and 15 m high in the canopy (“canopy traps”). At each stratum, traps were placed at three distances from the riverbank: 0 m (river edge), 250 m, and 500 m. To minimize disturbance associated with repeated human access along the transect route, traps were positioned approximately 50 m away from the central transect line. The six trap locations were permanent sampling stations established during long-term ecological studies at the site. The central transect had previously been surveyed using GPS, and the 0 m, 250 m, and 500 m positions were permanently marked in the field. Accordingly, the same tagged trees were used repeatedly throughout the present study rather than recording GPS coordinates for each sampling occasion. Traps were set between 17:00 and 17:30 and retrieved between 07:00 and 07:30 the following morning.

**Figure 1.**
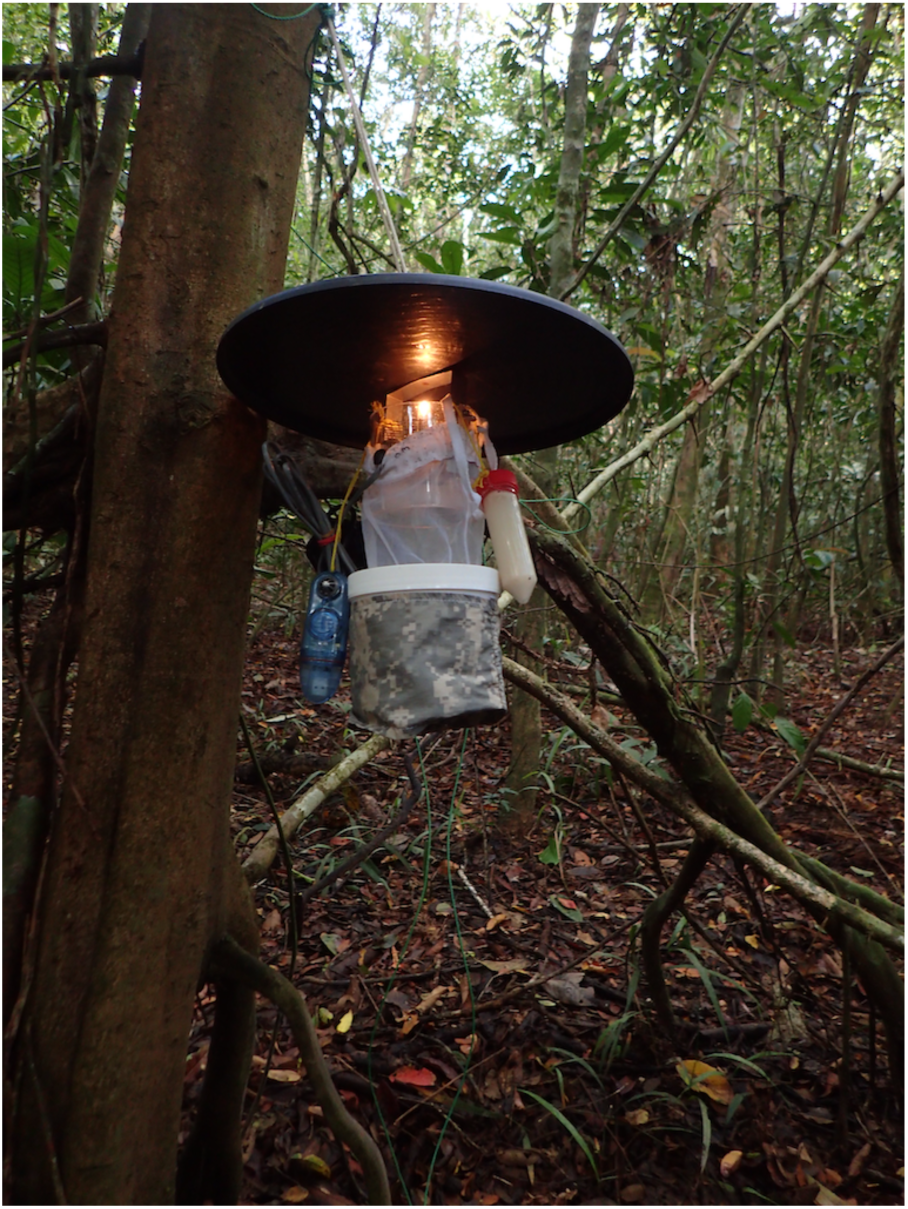
Mosquito trapping device used in the study. Each trap consisted of a battery-powered fan-based suction unit, a miniature light source, and a CO₂ generator (yeast–sugar mixture). Traps were installed either at ground level (1.5 m) or in the canopy (15 m) at distances of 0 m, 250 m, and 500 m from the Menanggul River.

### Mosquito identification and DNA extraction

Collected mosquitoes were transported to the field station and stored at −20 °C prior to processing. Specimens were morphologically identified to genus and, where possible, to species using standard taxonomic keys for Southeast Asian mosquitoes (35–38). Because our analyses focused on potential monkey malaria vectors, particular attention was given to *Anopheles* species in the Leucosphyrus Group, which were identified using their diagnostic morphological characters.

After identification, individual mosquitoes were preserved in 95% ethanol until DNA extraction. Only female *Anopheles* mosquitoes were selected for molecular screening, as males do not participate in pathogen transmission. DNA extraction and molecular screening for *Plasmodium* infections were conducted following previously published protocols developed in the laboratory of Tock-Hing Chua (24, 25). Briefly, genomic DNA was extracted from whole bodies of individual mosquitoes and initially screened for the presence of *Plasmodium* genus DNA using nested PCR assays targeting the small subunit ribosomal RNA (SSU rRNA) gene of *Plasmodium*. Samples that tested positive were subsequently subjected to species-specific PCR assays targeting simian malaria parasites, including *Plasmodium knowlesi*, *P. cynomolgi*, *P. inui*, *P. coatneyi*, and *P. fieldi*.

### Statistical analyses

Only female mosquitoes were included in the statistical analyses of overall mosquito abundance, because females are the blood-feeding and pathogen-transmitting sex. Male mosquitoes were recorded but excluded because of their limited relevance to biting risk and pathogen transmission. To visualize temporal trends in mosquito abundance, we first plotted the total number of female mosquitoes collected at each trap across time. Because log-transformation cannot be applied to zero values, observations with zero mosquitoes were assigned a value of 1 for visualization purposes only; all statistical analyses used the original untransformed counts. Temporal patterns were displayed using scatterplots with locally estimated scatterplot smoothing (LOESS), faceted by distance from the river (0 m, 250 m, 500 m) and sampling height (Ground or Tree). To evaluate spatial patterns in mosquito abundance, we fitted zero-inflated negative binomial generalized linear mixed models (ZINB GLMMs) using the glmmTMB package (39). Poisson GLMMs showed substantial overdispersion and excess zeros, supporting the use of a zero-inflated negative binomial framework (nbinom2 parameterization). The primary model included distance from the river (Location), trap height (Height), and their interaction as fixed effects, with sampling date (Date) included as a random intercept to account for repeated measurements:

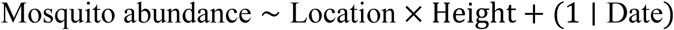

Because each distance × height combination was represented by a single trap on each sampling night, we could not include trap identity as a random effect. Instead, we accounted for temporal pseudoreplication by including sampling date as a random intercept, which treats observations from the same night as correlated. This approach is appropriate for designs without spatial replication at the trap level. To evaluate the robustness of the primary model to the influence of a single exceptionally large mosquito capture, we also conducted a sensitivity analysis excluding the trap collection with the highest mosquito count (400 individuals). For descriptive visualization of spatial differences, we generated log₁₀-scaled boxplots with jittered raw counts across Locations, again using 1 in place of zeros for plotting only. Estimated marginal means (EMMs) and corresponding confidence intervals were obtained from the ZINB GLMM using the *emmeans* package (40) to aid interpretation of model coefficients. In addition to the spatial model, we fitted a supplementary ZINB GLMM including night-time mean temperature and humidity as covariates. Environmental variables were derived from hourly logger data as nightly (17:00–07:00) means for each Location-Height combination and merged with trap counts; the nightly temperature and relative humidity data used in the analyses are provided in Table S1. Complete environmental records were available for 231 of the 244 trap collections, and the supplementary model including temperature and humidity was therefore fitted to this subset of the data. Model comparison using Akaike’s information criterion (AIC) was performed on models fitted to the same subset of data to evaluate whether inclusion of these covariates improved model performance. All analyses were conducted in R version 4.4.1 (41).

## Results

### Overall

Total mosquito abundance per sample ranged from 0 to 400 individuals (median = 30, IQR = 14–55), with substantial variation both temporally and across sites (Figure 2). Across all samples, a total of 9,928 mosquitoes were collected, and captures were strongly dominated by *Culex* spp. (9,079 individuals; 91.4% of all mosquitoes). By contrast, *Anopheles* spp. was rare (153 individuals; 1.5%), and only a single *Aedes* female was recorded. Because trapping was conducted overnight (17:00–07:30), when most *Aedes* species are relatively inactive, these collections were not designed to effectively sample diurnal mosquitoes. The remaining 695 mosquitoes (7.0%) comprised other mosquito taxa and specimens that could not be identified to genus because many were damaged and lacked the diagnostic morphological characters required for reliable identification. Sex ratios were markedly female-biased in the dominant genus (*Culex* spp.: 8,896 females vs. 183 males), consistent with the strong attraction of host-seeking females to light- and CO₂-baited traps. Zero-capture events occurred intermittently across locations and dates. Several distinct peaks in mosquito abundance were observed, with the largest spike occurring in July 2018 at the 500 m Tree site, where 400 individuals were captured in a single sample. Overall, the dataset reveals strong temporal fluctuations in mosquito abundance, with *Culex* spp. accounting for over 90% of all mosquitoes collected.

**Figure 2.**
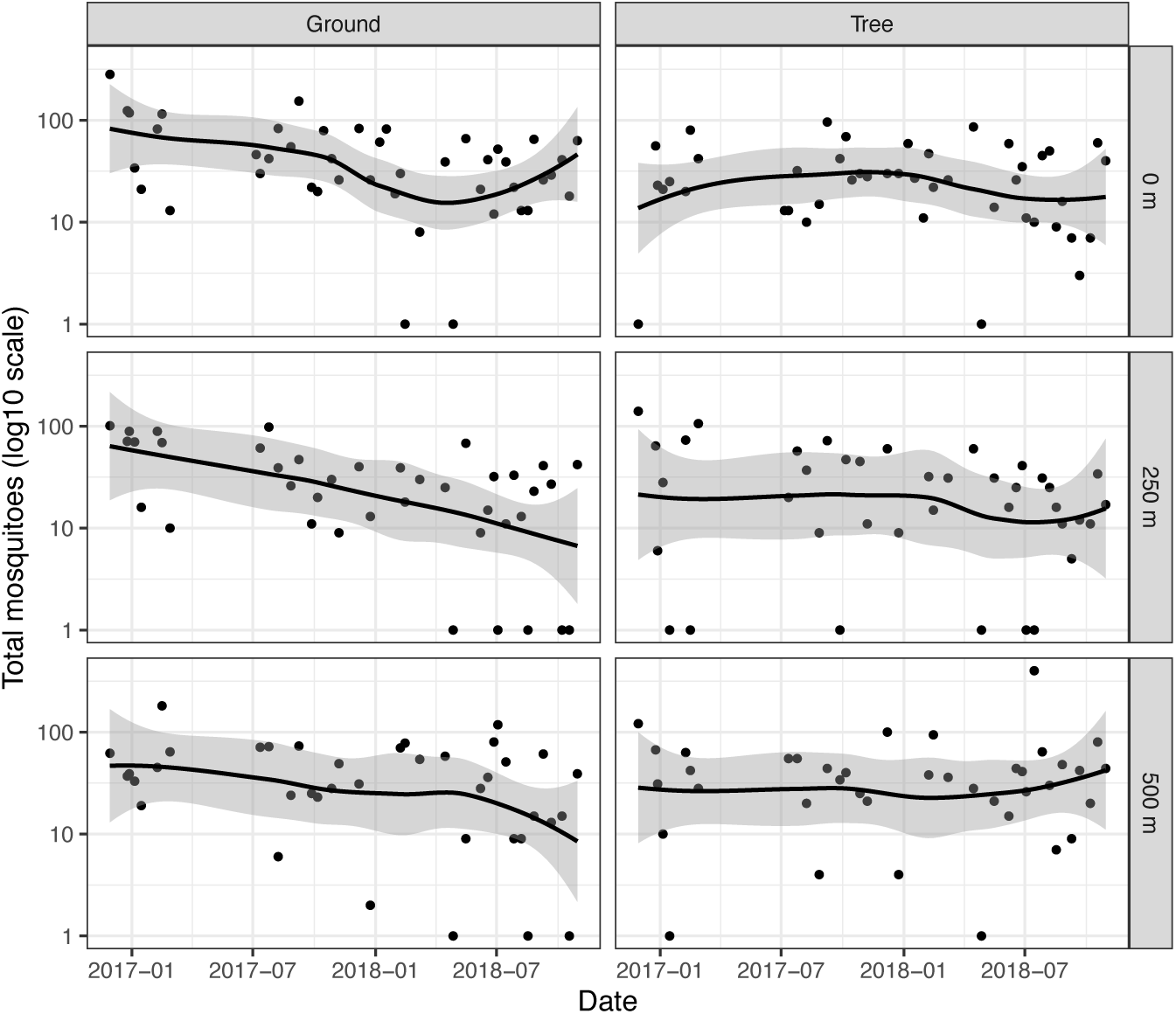
Temporal variation in female mosquito abundance from November 2016 to October 2018 across three distances from the river (0 m, 250 m, 500 m) and two sampling heights (Ground, Tree). Points represent the total number of female mosquitoes captured per sampling event, and lines show LOESS-smoothed trends with 95% confidence intervals (log₁₀ scale).

**Figure 3.**
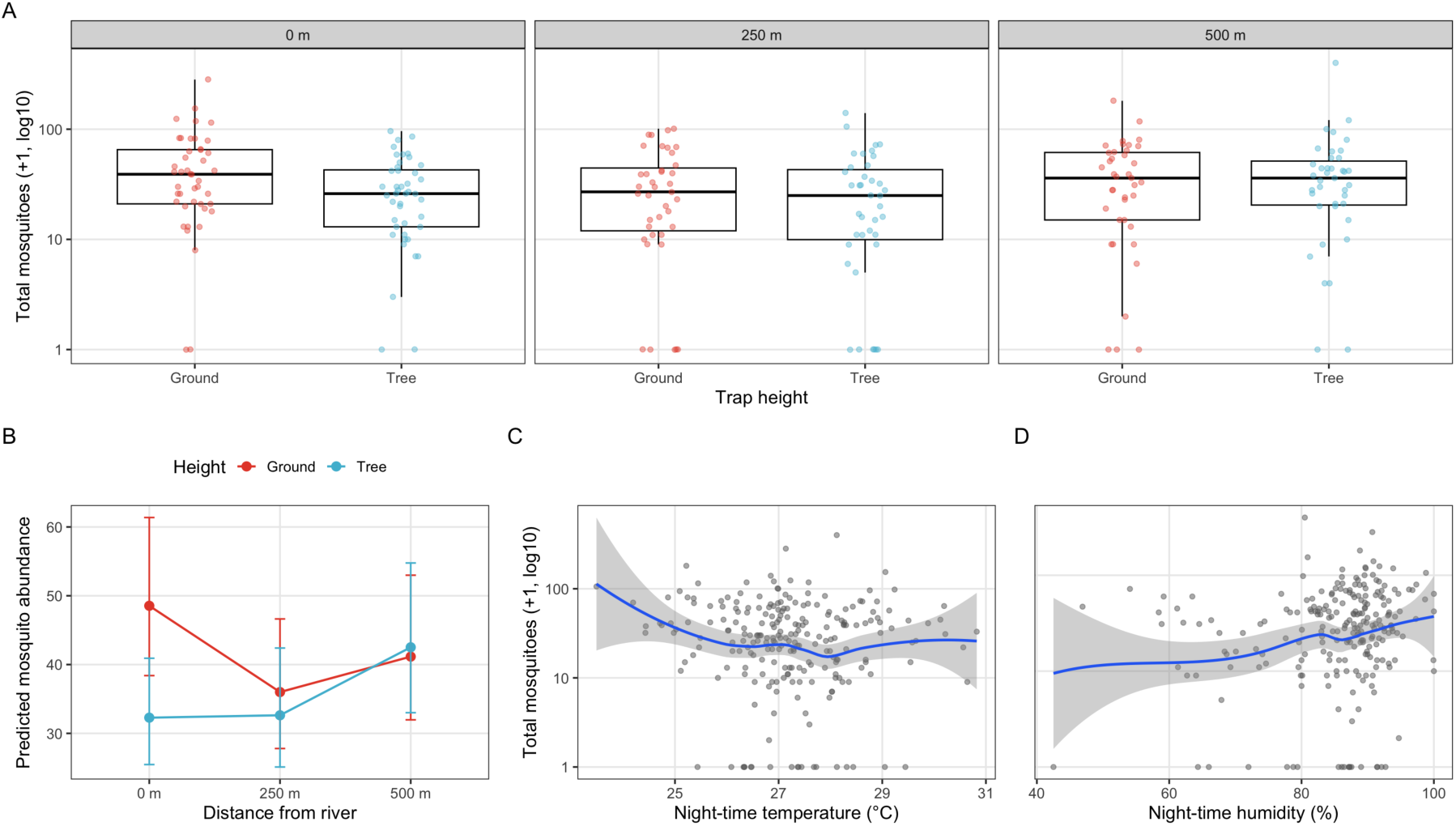
Spatial and microclimatic patterns in female mosquito abundance. (A) Differences in mosquito abundance across trap heights (Ground vs. Tree) and distances from the river (0 m, 250 m, 500 m). Boxplots show medians and interquartile ranges, with jittered points representing individual trap-night samples. Raw observations are shown, including repeated nightly measurements from the same traps. Mosquito counts are shown on a log₁₀ scale with zeros plotted as 1; (B) Model-predicted mosquito abundance from a zero-inflated negative binomial generalized linear mixed model including trap height, distance from the river, and their interaction, with sampling date included as a random effect. Points show estimated marginal means and error bars represent 95% confidence intervals; (C) Relationship between night-time mean temperature and mosquito abundance; (D) Relationship between night-time mean humidity and mosquito abundance. Panels C and D show samples matched to nightly logger data after quality control, with LOESS smoothers illustrating overall trends. Mosquito counts are shown on a log₁₀ scale for visualization.

### Effects of spatial factors on female mosquito abundance

A zero-inflated negative binomial GLMM revealed significant effects of distance from the river (Location) and sampling height (Height) on total mosquito abundance (Table 1). Mosquito counts tended to be lower at 250 m than at 0 m (β = –0.30, SE = 0.15, P = 0.049), corresponding to an approximately 26% decrease in expected abundance (Supplementary Table S2). Counts at 500 m did not differ significantly from those at the river edge (β = –0.17, P = 0.274). Sampling height also had a significant effect, with tree traps capturing fewer mosquitoes than ground traps (β = –0.41, SE = 0.14, P = 0.0047), corresponding to an approximately 34% reduction in abundance (Supplementary Table S2). In addition, the interaction between 500 m and tree height was significant (β = 0.44, SE = 0.21, P = 0.038), indicating that vertical differences in mosquito abundance varied across the river-distance gradient. However, sensitivity analysis excluding the single trap collection with the highest mosquito count (400 individuals) indicated that this interaction was no longer statistically significant (β = 0.29, SE = 0.20, P = 0.142), whereas the main effects of river distance and sampling height remained qualitatively unchanged. Collectively, these results indicate that mosquito abundance tends to be lower at intermediate distances from the river and remains consistently higher at ground level across the study area.

**Table 1.** Summary of zero-inflated negative binomial GLMM results testing the effects of distance from the river (Location), sampling height (Height), and their interaction on total mosquito abundance. The reference level is 0 m (Location), Ground (Height). Estimates are on the log scale; SE indicates standard error.

| Predictor | Estimate | SE | z-value | p-value |
| --- | --- | --- | --- | --- |
| Intercept (0m, Ground) | 3.8824 | 0.1196 | 32.45 | <0.001 |
| Location 250m | -0.2989 | 0.1516 | -1.97 | 0.0487 |
| Location 500m | -0.1650 | 0.1508 | -1.09 | 0.2737 |
| Height Tree | -0.4080 | 0.1442 | -2.83 | 0.0047 |
| 250m × Tree | 0.3096 | 0.2146 | 1.44 | 0.1491 |
| 500m × Tree | 0.4404 | 0.2124 | 2.07 | 0.0382 |

### Spatial patterns in potential monkey malaria vectors (*Anopheles*)

Female *Anopheles* mosquitoes were detected only sporadically in the trapping dataset (57 individuals in total). They were detected in only 36 of the 244 trap-nights (14.8%), further illustrating their rarity within the sampled mosquito community. Nevertheless, their occurrence was clearly structured across space and vertical strata (Table 2). Captures were concentrated near the river and were more frequent in ground traps than in tree traps. Over half of all female *Anopheles* were collected at 0 m (31/57, 54%), and ground traps captured more individuals than tree traps across the study area (32 vs. 25). Inland sites (250–500 m) yielded fewer captures overall (26/57, 46%), with particularly low numbers in inland canopy traps.

**Table 2.** Spatial and vertical distribution of *Anopheles* mosquitoes collected along the Menanggul River, Sabah, Malaysia. Counts are shown by species across distance from the river (0 m, 250 m, 500 m) and trap height (ground, tree). The total number of individuals per species and the number of *Anopheles* specimens positive for simian *Plasmodium* by PCR are also indicated.

| *Anopheles* spp. | Location & Place |  |  |  |  |  | Total | No. of *Anopheles* positive for malaria |
| --- | --- | --- | --- | --- | --- | --- | --- | --- |
|  | 0 |  | 250 |  | 500 |  |  |  |
|  | ground | tree | ground | tree | ground | tree |  |  |
| *An. balabacensis* | 10 | 14 | 4 | 2 | 4 | 1 | 35 | 2 |
| *An. donaldi* | 0 | 0 | 0 | 1 | 3 | 1 | 5 | 0 |
| *An. letifer* | 1 | 0 | 1 | 0 | 0 | 0 | 2 | 0 |
| *An. tessellatus* | 1 | 0 | 0 | 0 | 0 | 0 | 1 | 0 |
| *An. umbrosus* | 0 | 1 | 1 | 0 | 0 | 1 | 3 | 0 |
| *An. whartoni* | 1 | 0 | 0 | 1 | 1 | 0 | 3 | 0 |
| *unidentified* | 2 | 1 | 3 | 2 | 0 | 0 | 8 | 0 |
|  | 15 | 16 | 9 | 6 | 8 | 3 | 57 | 2 |

### Species composition of *Anopheles* mosquitoes

A total of 57 female *Anopheles* mosquitoes were collected, representing six identified species plus unidentified specimens (Table 2). The assemblage was dominated by *An. balabacensis* (35/57; 61%), which occurred across all distances from the river. The remaining species (*An. donaldi*, *An. umbrosus*, *An. whartoni*, *An. letifer,* and *An. tessellatus*) were detected only sporadically and at low frequencies.

### Detection of monkey *Plasmodium* in *Anopheles* mosquitoes

PCR screening detected *Plasmodium* DNA in two *Anopheles* specimens, both identified as *An. balabacensis* (Table 2). No *Plasmodium* DNA was detected in any other *Anopheles* species. Mosquitoes were screened for multiple simian *Plasmodium* species, including *P. coatneyi*, *P. inui*, *P. fieldi*, *P. cynomolgi,* and *P. knowlesi*. Species-specific PCR indicated mixed infections in both positive mosquitoes: one individual was positive for *P. coatneyi*, *P. fieldi*, and *P. inui*, whereas the other was positive for *P. coatneyi*, *P. cynomolgi*, *P. inui*, and *P. knowlesi*. These results demonstrate the occurrence of *Plasmodium* DNA-positive *An. balabacensis* within the trapping area.

### Effects of microclimatic conditions on mosquito abundance

To assess whether nightly microclimatic variation contributed additional explanatory power beyond spatial factors, we fitted a supplementary zero-inflated negative binomial model including night-time mean temperature and humidity. Neither temperature (β = – 0.09, SE = 0.07, P = 0.185) nor humidity (β = 0.003, SE = 0.009, P = 0.750) significantly predicted mosquito abundance (Table S3). Model comparison using AIC indicated little difference between the spatial model and the model including microclimatic covariates fitted to the same subset of data (AIC = 2101.0 vs. 2099.8; ΔAIC = 1.23), suggesting that the inclusion of temperature and humidity did not substantially improve model fit. These results indicate no statistically significant effects of night-time temperature or humidity on mosquito abundance within the range of environmental conditions sampled.

## Discussion

This study provides an integrated assessment of mosquito community structure and fine-scale spatial distribution in a tropical riverine forest in Sabah, Malaysian Borneo. Mosquito assemblages were strongly dominated by *Culex* spp., while *Anopheles*, including potential monkey malaria vectors, were detected only sporadically. Nevertheless, female *Anopheles*, particularly *An. balabacensis*, showed suggestive spatial patterns associated with river proximity and ground-level traps, and included *Plasmodium* DNA-positive individuals, underscoring their potential epidemiological relevance. Together, these findings suggest that vector abundance alone may be a poor proxy for transmission risk; instead, zoonotic risk is likely to be shaped by the spatial overlap among vectors, primate hosts, and human activity in tropical landscapes, which jointly determines exposure at forest-human interfaces (16, 18, 24).

Due to the low number of *Anopheles* captures (n = 57) and the lack of trap replication, we did not attempt formal statistical modeling of their spatial distribution. Instead, the patterns described here should be interpreted as descriptive observations from this specific site rather than statistically robust inferences. Future studies with replicated sampling designs across multiple riverine systems will be required to evaluate the generality of these patterns more rigorously.

### Vector community in the Borneo context

The low relative abundance of *Anopheles* in our collections may partly reflect differences in collection efficiency among mosquito taxa when using light-and CO₂-baited traps, which do not necessarily track host-vector contact rates directly (8, 42). Accordingly, trap-based estimates should be interpreted as indices of host-seeking activity rather than direct measures of biting exposure. Recent work in monkey malaria systems further emphasizes that transmission risk is shaped not only by vector abundance but also by the spatial overlap among vectors, reservoir hosts, and human activities across forest, farm, and village interfaces (16, 18, 24).

Importantly, despite being rare in the overall assemblage, *An. balabacensis* is widely recognized as a primary vector of *Plasmodium knowlesi* and other monkey malaria parasites in Malaysian Borneo, and infected mosquitoes have been detected in forest and forest-edge settings in Sabah (13, 14). The dominance of *An. balabacensis* among the *Anopheles* captured in our study is therefore consistent with regional patterns, underscoring that epidemiologically important vectors may contribute disproportionately to zoonotic transmission even when they comprise only a small fraction of the mosquito community. Longitudinal studies in Sabah have shown that the infection prevalence of *An. balabacensis* with simian *Plasmodium* is typically low, yet this species is consistently identified as the primary vector of *P. knowlesi*, with parasite lineages matching those found in both humans and macaques (14, 25). Together with our detection of simian *Plasmodium* DNA in *An. balabacensis*, this supports the view that a small number of highly competent vectors in ecologically strategic microhabitats can sustain appreciable zoonotic risk.

### Environmental and spatial drivers of mosquito abundance

Mosquito abundance was significantly structured by space, with lower counts at intermediate distances from the river (250 m), while no clear difference was observed between the river edge (0 m) and the furthest distance (500 m). In addition, mosquito captures were consistently lower in canopy traps. Furthermore, the significant interaction between distance and height suggests that vertical differences in mosquito abundance varied across the river-distance gradient, indicating that canopy–ground contrasts are not uniform across space. Because the assemblage was overwhelmingly dominated by *Culex*, these patterns likely reflect the spatial ecology of the dominant taxon, whose adult distributions are shaped by the availability and configuration of aquatic breeding sites and associated habitat features (8, 13, 20). Recent analyses of *An. balabacensis* larval habitats in Sabah indicate that breeding sites are more frequently associated with fragmented forest edges and certain plantation types, particularly oil palm and rubber, than with undisturbed primary forest (20). These findings, together with our observation that adult mosquito abundance was structured by distance to the river and vertical stratum, suggest that spatial heterogeneity in surrounding landscape structure may influence both larval production and adult host-seeking activity in riverine systems. The lower abundance observed in canopy traps is also consistent with vertical stratification commonly reported for forest mosquitoes, which can arise from height-specific host availability, flight behaviour, and microhabitat conditions (8, 9).

A strength of our study design was the deployment of temperature and humidity loggers at each trap, which allowed us to link nightly microclimatic conditions to mosquito captures while accounting for site- and date-specific variation. Within the range of temperatures and humidities sampled, however, we detected no clear association between night-time mean conditions and mosquito abundance. Although microclimate can influence mosquito development and survivorship in other contexts (6, 7), no statistically significant effects of short-term temperature or humidity were detected within the environmental range sampled in this study.

Temporal patterns in trap captures also suggested that mosquito abundance was not synchronized across sites. For example, a pronounced peak was observed in the 500 m canopy trap during July 2018, whereas river-edge traps showed relatively more stable captures through time. This pattern may reflect localized environmental conditions or stochastic variation in breeding-site dynamics rather than uniform seasonal fluctuations across the forest, although our sampling was not designed to test seasonality explicitly. To evaluate whether this single exceptionally large capture disproportionately influenced the GLMM results, we conducted a sensitivity analysis excluding this observation. The main effects of river distance and sampling height remained qualitatively unchanged, whereas the interaction between river distance and sampling height was no longer statistically significant. These findings indicate that the main spatial patterns were robust, whereas evidence for the interaction should be interpreted cautiously at this point.

Overall, these results highlight pronounced horizontal and vertical heterogeneity in mosquito abundance across the forest, which may generate localized hotspots of vector-host contact depending on where hosts and humans concentrate their activities. The use of zero-inflated models indicates that the observed mosquito counts included more zero captures than expected under a standard negative binomial model, consistent with pronounced spatial heterogeneity in mosquito occurrence.

### Spatial ecology of potential vectors

Although *Anopheles* mosquitoes were rare overall, their capture showed suggestive spatial structuring. Captures were most frequent at the river edge (0 m). *Anopheles* were collected across both vertical strata, with a modest overall tendency for more captures in ground traps than in those placed higher up in the canopy, suggesting that potential vectors may be concentrated near riverine habitats while not being strictly confined to a single forest stratum. This pattern is broadly consistent with mosquito life history: immature stages depend on aquatic breeding sites, and adult distributions can be shaped by the availability and spatial arrangement of those habitats (2, 8). In addition, *Anopheles* species often exhibit host species-specific preferences and host-seeking behaviour, which can influence spatial patterns of host-vector contact (2, 8). Even with the limitations of trap-based sampling, the higher capture frequency of *Anopheles* at the river edge in this study suggests that river-edge habitat may represent localized zones of increased vector occurrence, particularly when host activity overlaps temporally with vector host-seeking (14, 17).

Importantly, however, these inferences are based on light- and CO₂-baited trap collections, and trap catches do not necessarily translate directly into realized biting rates or host–vector contact. Differences among collection methods can yield markedly different estimates of vector abundance and composition, and trap-based indices may under- or over-represent exposure to particular taxa (42–44). Therefore, the lower relative abundance of *Anopheles* in canopy traps should not be interpreted as definitive evidence of negligible biting exposure in the strata. This pattern may also partly reflect differences in trap detectability within structurally complex canopy environments, where CO₂ plumes and light cues may disperse differently than near the forest floor.

### Zoonotic risk and human–wildlife interfaces

Zoonotic risk is expected to be highest where competent vectors, reservoir hosts, and human activity overlap in space and time. In our study, *Anopheles* mosquitoes were rare but spatially structured, being recorded more often near the river and primarily in ground traps. Given the established role of *Anopheles balabacensis* as a primary vector of simian malaria parasites in Sabah (13–15), the detection of *Plasmodium* DNA in two *An. balabacensis* individuals, including one positive for *P. knowlesi* together with other simian *Plasmodium*, species suggests that *Plasmodium* transmission may be occurring within the study area.

We detected *Plasmodium* DNA in two *An. balabacensis* individuals (one from 0 m ground trap, one from 0 m canopy trap). While this demonstrates that *Plasmodium*-positive *An. balabacensis* were present at the study site, the small number of positives precludes any inference about spatial patterns of infection risk or true prevalence. These detections should be interpreted as qualitative evidence of local *Plasmodium* circulation rather than quantitative estimates of infection pressure. Because PCR was performed on whole mosquito bodies, the detected *Plasmodium* DNA may have originated from either infected blood meals or infective sporozoites in the salivary glands. Therefore, these results should not be interpreted as direct evidence of vector competence.

At this specific site along the Menanggul River, the river edge showed higher mosquito capture rates and greater *Anopheles* presence than inland locations. If this pattern holds more broadly, riverine forests could represent hotspots of vector-host contact, but replicated studies across multiple river systems are needed to test this hypothesis. In general, macaques are widely implicated as natural hosts for diverse *Plasmodium* taxa across Asia (23). Indeed, in the Lower Kinabatangan region, macaques are recognized as reservoir hosts of simian malaria parasites (24). In this landscape, several primate species—including long-tailed and pig-tailed macaques as well as proboscis monkeys—frequently use river-edge habitats, particularly at dawn and dusk when they are most readily observed (29). Because these periods often coincide with peak activity of many mosquito species, they may increase opportunities for host–vector contact.

Human activity may further amplify this spatial and temporal overlap. The Menanggul River is a well-known ecotourism site where visitors commonly observe primates from boats at dawn and dusk, resulting in repeated human presence in river-edge habitats during periods of high mosquito activity (27, 28). Indeed, human-landing studies in Sabah further show that *An. balabacensis* tends to bite predominantly at ground level and exhibits early evening biting peaks (14), reinforcing that periods when primates and humans use river-edge habitats may coincide with elevated vector activity. Such temporal and vertical alignment likely amplifies opportunities for host-vector contact in these microhabitats (19).

Previous studies have also shown that *P. knowlesi* transmission is closely associated with human-wildlife interfaces, where forest use, agricultural activities, and ecotourism bring humans into contact with both reservoir hosts and vector mosquitoes (15–18, 21, 22). Population-based, case-control, and sero-epidemiological studies in Sabah further demonstrate that overnight stays in or near forests, recent forest visits, and work in plantations adjacent to forest edges are associated with increased odds of *P. knowlesi* infection (15, 21). In riverine ecotourism settings such as the Kinabatangan, repeated dawn and dusk primate-viewing activities may therefore place visitors within these high-risk interfaces at times that overlap with peak *An. balabacensis* biting, potentially increasing spillover opportunities even when vector densities are low (14). Under such conditions, even relatively low vector densities may be sufficient to sustain spillover transmission if spatial and temporal overlap is high.

Several limitations should be considered when interpreting these findings. Most importantly, our study lacks true replication at the trap level, with only one trap per distance × height combination. This design choice, necessitated by logistical constraints associated with canopy access in a remote field site, means that our inferences about spatial patterns are strictly applicable to these specific sampling locations along a single transect. We cannot formally separate the effects of distance from the river or vertical stratum from site-specific characteristics of individual trap locations. Consequently, the apparent effects of river distance and vertical stratum should be interpreted cautiously, as they may partly reflect characteristics unique to the individual trap locations rather than these spatial factors themselves. Therefore, while our results document spatial variation at this site, they are best interpreted as hypotheses about general patterns in riverine forests rather than definitive tests. Future studies with replicated traps across multiple riverine systems will be necessary to establish the generality of these patterns.

Taken together, our findings are consistent with the emerging consensus that zoonotic malaria risk in Sabah is driven less by overall *Anopheles* abundance and more by localized “contact hotspots” where competent vectors, macaque and possibly langur reservoirs, and humans repeatedly co-occur in space and time. Riverine forest edges that attract both wildlife and people, particularly in fragmented and agriculturally modified landscapes, may therefore represent plausible foci for *P. knowlesi* spillover, despite low trap-based estimates of vector density. Future efforts should also focus on detecting malaria infections in the local primate population.

These findings also have potential implications for public health and ecotourism management in riverine landscapes. If zoonotic malaria risk is concentrated within localized contact hotspots, interventions may be more effective when targeted toward high-risk microhabitats and periods of peak human-vector overlap rather than applied uniformly across broader forest landscapes. In systems such as the Kinabatangan, where ecotourism activities frequently occur at dawn and dusk along river edges, practical measures including the use of topical repellents, long-sleeved clothing, and increased awareness of mosquito exposure may help reduce opportunities for spillover transmission. More broadly, our findings support the growing recognition that land-use change, riverine habitat structure, wildlife behaviour, and human activities should be considered together within an integrated One Health framework for managing zoonotic disease risk in tropical landscapes.

## Acknowledgements

We express our sincere thanks to the Sabah Biodiversity Centre and the Sabah Wildlife Department for granting permission to carry out this research. We are also deeply indebted to the Sabah Forestry Department for facilitating the use of their facilities in the field. We particularly thank the research assistants, Asnih Binti Etin and Jasrudy Bin Mandu, for their support in the field. We also acknowledge the Collaborative Research Program of the Wildlife Research Center, Kyoto University, for facilitating academic exchange related to this study.

## Author contribution

IM: Writing – original draft, Visualization, Formal analysis, Field investigation, Methodology, Project administration, Conceptualization; BOM: Mosquito identification, Molecular analyses, Data curation, Writing – review & editing; TY: Writing – review & editing, Discussion of zoonotic disease ecology; PL: Writing – review & editing, Resources; JT: Writing – review & editing, Resources; MAH: Writing – review & editing, Discussion of monkey malaria ecology, Resources; HB: Writing – review & editing, Discussion from primate ecological perspectives, Resources; VSK: Writing – review & editing, Discussion of molecular and zoonotic disease aspects; THC: Supervision of mosquito identification and molecular analyses, Resources, Writing – review & editing.

## Funding

This study was supported by Japan Society for the Promotion of Science KAKENHI (24H00774), the Core-to-Core Program, Asia-Africa Science Platforms (JPJSCCB20250006), the Research Center for GLOBAL and LOCAL Infectious Diseases, Oita University (2024B12), and the AMED (JP26jm0110032). The funders had no role in study design, data collection and analysis, decision to publish, or preparation of the manuscript.

## Data Availability

All relevant data underlying the findings of this study are provided within the manuscript and its Supporting Information files.

## Declarations

### Ethics approval

Research permits for this study were granted by the Sabah Biodiversity Centre and the Sabah Wildlife Department [JKM/MBS.100-2/2 JLD.5 (76)].

### Conflict of interest

The authors have declared that no competing interests exist.

### Generative AI and AI-assisted technologies in the writing process

During the preparation of this manuscript, the authors used ChatGPT (OpenAI) to refine the clarity and logical flow of the text. The authors carefully reviewed, corrected, and approved all content generated, and take full responsibility for the final published version.

## Notes

### Competing Interest Statement

The authors have declared no competing interest.

### Summary of Updates

We improved the text based on the reviewers' comments.

