## Supplemental Files for "Spatial patterns of mosquito communities and monkey malaria vectors in a tropical riverine forest in Sabah, Malaysian Borneo" (from another version of this preprint: version 1, 2026-04-28)

Supplementary Table 1S **Effect sizes from the zero-inflated negative binomial GLMM examining the influence of distance from the river (Location), trap height (Height), and their interaction on total mosquito abundance.** Values represent exponentiated coefficients (exp(Estimate)), interpreted as multiplicative changes in expected mosquito counts relative to the reference category (0 m, Ground). Shown are point estimates and corresponding 95% confidence intervals.

| Predictor | exp(Estimate) | 95% CI |
| --- | --- | --- |
| Location 250m | 0.74 | 0.55–1.00 |
| Location 500m | 0.85 | 0.63–1.14 |
| Height (Tree) | 0.66 | 0.50–0.88 |
| 250m × Tree | 1.36 | 0.89–2.08 |
| 500m × Tree | 1.55 | 1.02–2.36 |

Supplementary Table 2S Parameter estimates and exponentiated effect sizes from the zero-inflated negative binomial GLMM evaluating the influence of night-time temperature and humidity on mosquito abundance. The model included distance from the river (Location), sampling height (Height), and their interaction as fixed effects, and sampling date as a random intercept. Estimates are presented on the log scale, along with standard errors (SE), z-values, and p-values. Exponentiated coefficients (exp(Estimate)) and 95% confidence intervals represent multiplicative changes in expected mosquito counts relative to the reference category (0 m, Ground). Neither temperature nor humidity significantly predicted mosquito abundance, and inclusion of these covariates did not improve model performance relative to the spatial model.

| **Predictor** | **Estimate** | **SE** | **z-value** | **p-value** | **exp(Estimate)** | **95% CI** |
| --- | --- | --- | --- | --- | --- | --- |
| Night-time temperature | -0.09434 | 0.07117 | -1.325 | 0.185 | 0.91 | 0.79–1.05 |
| Night-time humidity | 0.00287 | 0.00898 | 0.319 | 0.75 | 1 | 0.98–1.02 |
| Location 250 m | -0.3094 | 0.15936 | -1.941 | 0.052 | 0.73 | 0.53–1.00 |
| Location 500 m | -0.22616 | 0.16327 | -1.385 | 0.166 | 0.8 | 0.58–1.12 |
| Height (Tree) | -0.47612 | 0.16263 | -2.928 | 0.003 | 0.62 | 0.45–0.86 |
| 250 m × Tree | 0.38996 | 0.28289 | 1.379 | 0.168 | 1.48 | 0.85–2.58 |
| 500 m × Tree | 0.53134 | 0.23637 | 2.248 | 0.025 | 1.7 | 1.07–2.70 |
